# *De novo* phased assembly of the *Vitis riparia* grape genome

**DOI:** 10.1101/640565

**Authors:** Nabil Girollet, Bernadette Rubio, Pierre-François Bert

**Author notes:** Corresponding Author: Pierre-François Bert.

## Abstract

Grapevine is one of the most important fruit species in the world. In order to better understand genetic basis of traits variation and facilitate the breeding of new genotypes, we sequenced, assembled, and annotated the genome of the American native *Vitis riparia*, one of the main species used worldwide for rootstock and scion breeding. A total of 164 Gb raw DNA reads were obtained from *Vitis riparia* resulting in a 225X depth of coverage. We generated a genome assembly of the *V. riparia* grape *de novo* using the PacBio long-reads that was phased with the 10x Genomics Chromium linked-reads. At the chromosome level, a 500 Mb genome was generated with a scaffold N50 size of 1 Mb. More than 34% of the whole genome were identified as repeat sequences, and 37,207 protein-coding genes were predicted. This genome assembly sets the stage for comparative genomic analysis of the diversification and adaptation of grapevine and will provide a solid resource for further genetic analysis and breeding of this economically important species.

## Background & summary

Since few decades and the development of sequencing technologies, the number of species whose genome has been totally sequenced has increased exponentially. There is a large variability for the quality of all the sequences assemblies. In 2017, 72 plant reference quality genome assemblies were reported in NCBI^1^. For plant breeding, the availability of a contiguous genome sequence provides a tool to better identify genes underlying traits and how they may be regulated by various environmental parameters in different genetic backgrounds. At the simplest, it allows for association of genetic markers for selection and introgression of traits across germplasm to enable the development of novel products for consumers^2 3^.

As an important crop, *Vitis vinifera* was one of the first higher plant species whose genome was sequenced by a French-Italian consortium^4^. The consortium decided to sequence a near homozygous *V. vinifera* cultivar related to Pinot Noir (PN40024) in order to facilitate the sequence assembly by limiting sequence variability. To date, this genome still stands as the reference for the grapevine community, but grapevine intra species and interspecies diversity makes using a single reference genome inadequate for studying the function of other genotypes. In order to address the variations in a cultivated *V. vinifera* variety, the Pinot Noir genome was sequenced using Sanger sequencing providing a high quality draft of the genome with about 10X coverage^5^. Next Generation Sequencing reads are too short to resolve abundant repeats in particular in plants genome, leading to incomplete or ambiguous assemblies^6^. Few attempts to produce high quality grapevine genomes were undertaken in grapevine and produced valuable data to study the genetic variations of *V. vinifera* cv. Tannat^7^ and cv. Thompson seedless^8^ through comparison with the reference genomes.

The last few years have seen rapid innovations in sequencing technologies and improvement in assembly algorithms that enabled the creation of highly contiguous genomes. The development of third generation sequencing technologies that deliver long reads from single molecules and carry the necessary information to phase haplotypes over several kilobases have greatly improved the feasibility of *de novo* assemblies^9 10 11^. Sequences of *V. vinifera* cv. Cabernet Sauvignon were first released^12^ using PacBio sequencing and FALCON, and FALCON-Unzip pipeline^12^. This generated a 591 Mbp haplotype genome from a set of 718 primary contigs, and a set of correlated 2,037 haplotigs spanning 367 Mbp. The total p-contig size was larger than the estimated genome size of *V. vinifera* (∼500 Mbp) suggesting that in some cases FALCON-Unzip underestimated the alternative haplotype sequences because of high heterozygosity between homologous regions, which is common in grapevine^13 14^. Later, the PacBio assembly and annotation of *V. vinifera* cv Chardonnay variety provided after curation of artefactual contig assignment, 854 p-tigs and 1883 h-tigs, totaling 490 Mb and 378 Mb ^15^. More recently, another version of the Chardonnay genome was proposed with a different level of curation at 605 Mb^16^.

An evaluation of genetic diversity based on a panel of 783 *V. vinifera* varieties using 10K SNPs revealed a high level of diversity (He = 0.32) and confirmed the close pedigree relationship within the cultivated grapevine due to the wide use of the most interesting parents during domestication and early selection by humans^17^. Considering that grape cultivation currently faces severe pathogen pressures and climate change, we assume that the exploitation of the natural genetic diversity may ensure the long-term sustainability of the grape and wine industries^18^. Grapes belong to the genus *Vitis*, which includes over 60 inter-fertile species. The most common grape cultivars derive their entire ancestry from the species *V. vinifera*, but wild relatives have also been exploited to create hybrid cultivars, often with increased disease resistances^19^.

To date, no wild *Vitis* genomes have been released so far and the only whole genome sequences for grape are from *V. vinifera* varieties and yet there is a clear need for genetic resources ^20^. Here, we report the first *de novo* assembly and genome annotation of the North American native grape *V. riparia*. Using the latest sequencing technologies, we show that 10x Genomics Chromium data can be combined with long read PacBio sequencing to effectively determine genome phasing. The phased haplotypes of *V. riparia* genome will greatly contribute to give more insight into the functional consequences of genetic variants.

## Methods

### Sample collection, library construction and whole genome sequencing

The *Vitis riparia* Gloire de Montpellier (RGM) selection was obtained in 1880 by L. Vialla and R. Michel from North American collections and is the only commercially available pure *V. riparia* stock. RGM clone #1030 and the European native *Vitis vinifera* Cabernet sauvignon (CS) clone #15 were grown at INRA, Bordeaux (France). A F1 segregating population of 114 individuals named CSxRGM1995-1 was derived from the cross between CS and RGM^21^. This population was genotyped using the GBS approach^22^ to create a high resolution genetic map to assist in anchoring and orienting the assembled *V. riparia* genome scaffolds.

Total DNA was isolated and extracted using QIAGEN Genomic-tips 100/G kit (Cat No./ID: 10243) following the tissue protocol extraction. Briefly, 1g of young leaf material was ground in liquid nitrogen with mortar and pestle. After 3h of lysis and one centrifugation step, the DNA was immobilized on the column. After several washing steps, DNA is eluted from the column, then desalted and concentrated by alcohol precipitation. The pellet is resuspended in TE buffer.

Three PacBio libraries with a 20-kb insert size were also constructed and sequenced on RSII platforms (97.71 Gb data; ∼118-fold covering), following the standard PacBio protocol of Sequencing Kit 1.2.1 (Pacific Biosciences, USA). Four 10x Chromium Genomics libraries were constructed using the Chromium™ Genome Solution (10X Genomics, USA), and 2×150 bp sequenced on Illumina HiSeq3000, producing ∼350 million paired-end linked-reads (∼ 107-fold covering). Finally, 2 libraries for 2×100 bp sequencing were built with different insert sizes: 500 bp for paired-end (PE) and 6 kb for mate-pair (MP), based on the standard Illumina protocol and sequenced on the Illumina HiSeq 2500. The raw reads were trimmed before being used for subsequent genome assembly. For Illumina HiSeq sequencing, the adaptor sequences, the reads containing more than 10% ambiguous nucleotides, as well as the reads containing more than 20% low-quality nucleotides (quality score less than 5), were all removed. After data cleaning and data preprocessing, we obtained a total of 164 Gb of clean data (52 Gb PacBio data, 59 Gb 10X Genomics, 33 Gb PE reads and 20 Gb MP reads,), representing 331X coverage of the *V. riparia* genome (Table 1).

**Table 1:**
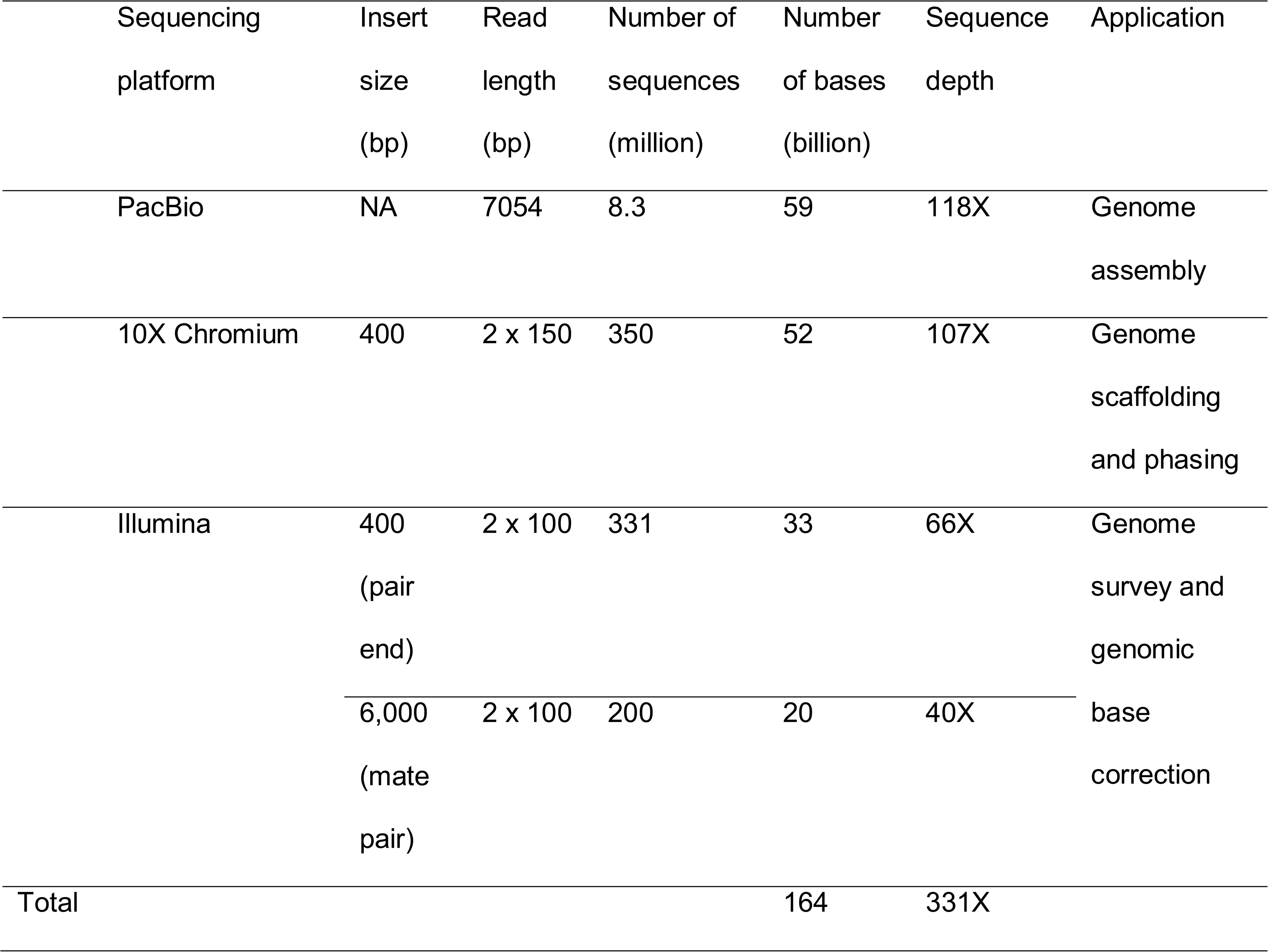
Data count and library informations for *Vitis riparia* genome sequencing.

### Genome size and heterozygosity estimation

Lodhi and Reisch^35^ estimated the genome size in grape to be approximately 475 Mb based on measurements using flow cytometry for 19 species including wild *Vitis* species, *V.vinifera* and *V. labrusca* cultivars. The measurements showed intraspecific variation in genome size between different varieties of *Vitis vinifera* ranging from 1C = 415 to 511 Mb, and between different North America *Vitis* species ranging from 1C = 411 to 541 Mb, with *V. riparia* around 470 Mb. Genome sequencing of different *V. vinifera* varieties gave values in the same range or greater depending on the methods of sequencing and assembly. In order to verify these values, we estimated genome size of *V. riparia* by the k-mer method^36 37^ using data from pair- end and mate-pair Illumina sequencing. By analyzing the 21-mers depth distribution, a total of ∼50 billion k-mers were estimated with a peak frequency of 100, corresponding to a genome size of 494 Mb and the estimated repeat sequencing ratio was 33.74%. In this study, *V. riparia* heterozygosity was estimated to be 0.46% (mean distance 1 SNP each 217 bp between heterozygous SNPs) from 10x Chromium Genomics data processing.

### *De novo* Genome assembly and scaffolding of the *Vitis riparia* genome

We employed a hybrid *de novo* whole-genome assembly strategy, combining both short linked-reads and PacBio long reads data. Genome assembly was first performed on full PacBio cleaned reads using FALCON v0.3.0^38^. Error correction and pre-assembly were carried out with the FALCON/FALCON Unzip pipeline after evaluating the outcomes of using different parameters in FALCON during the pre-assembly process. Based on the contig N50 results, a *length_cutoff* of 5kb and a *length_cutoff_pr* of 8kb for the assembly step were ultimately chosen. The draft assembly was polished using Quiver^39^, which mapped the PacBio reads to the assembled genome with the BLASER pipeline^40^. Haplotypes were separated during assembly using FALCON-Unzip and the preliminary genome assembly was approximately 530 Mb (1,964 primary-contigs) and 317 Mb (3,344 haplotigs). A summary of the assembly statistics can be found in Table 1. Assembly was then processed with Purge Haplotigs^13^ to investigate the proper assignment of contigs, followed by 2 rounds of polishing to correct residual SNP and INDELs errors with Pilon v1.22 software^41^ using high-coverage (∼106X) Illumina paired-end and mate pair data.

The 10x Chromium Genomics linked-reads were used to produce a separate *V. riparia* assembly using the Supernova assembler option *--style=pseudohap2* and created two parallel pseudohaplotypes^42^. The mean input DNA molecule length reported by the Supernova assembler was 45kb and the assembled genome size was 424Mb with a N50 scaffold of 711kb.

Subsequently, the PacBio assembly was scaffolded with the 10x Chromium Genomics one using the hybrid assembler LINKS^43^ with 7 iterations, producing 870 scaffolds spanning 500 Mb with N50 = 964 kb and L50 = 255 (Table 2). Finally, genome phasing was reconstituted using Long Ranger analysis pipeline that processes Chromium sequencing output to align reads and call and phase SNPs, indels, and structural variants on the basis of molecular barcodes information.

**Table 2:**
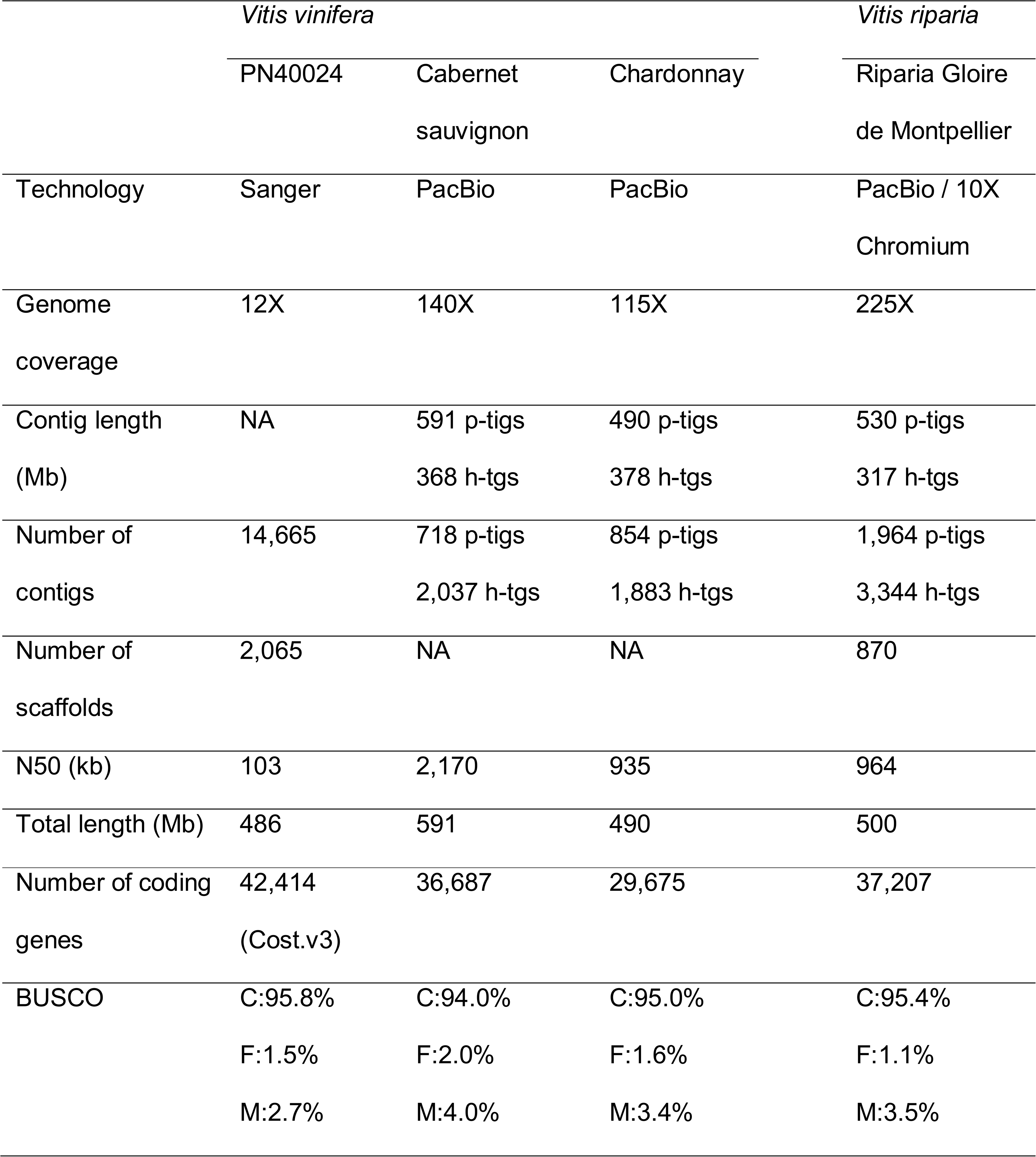
Summary of the *V. riparia* genome assembly and comparison with with *V. vinifera* varieties.

**Table 3:**
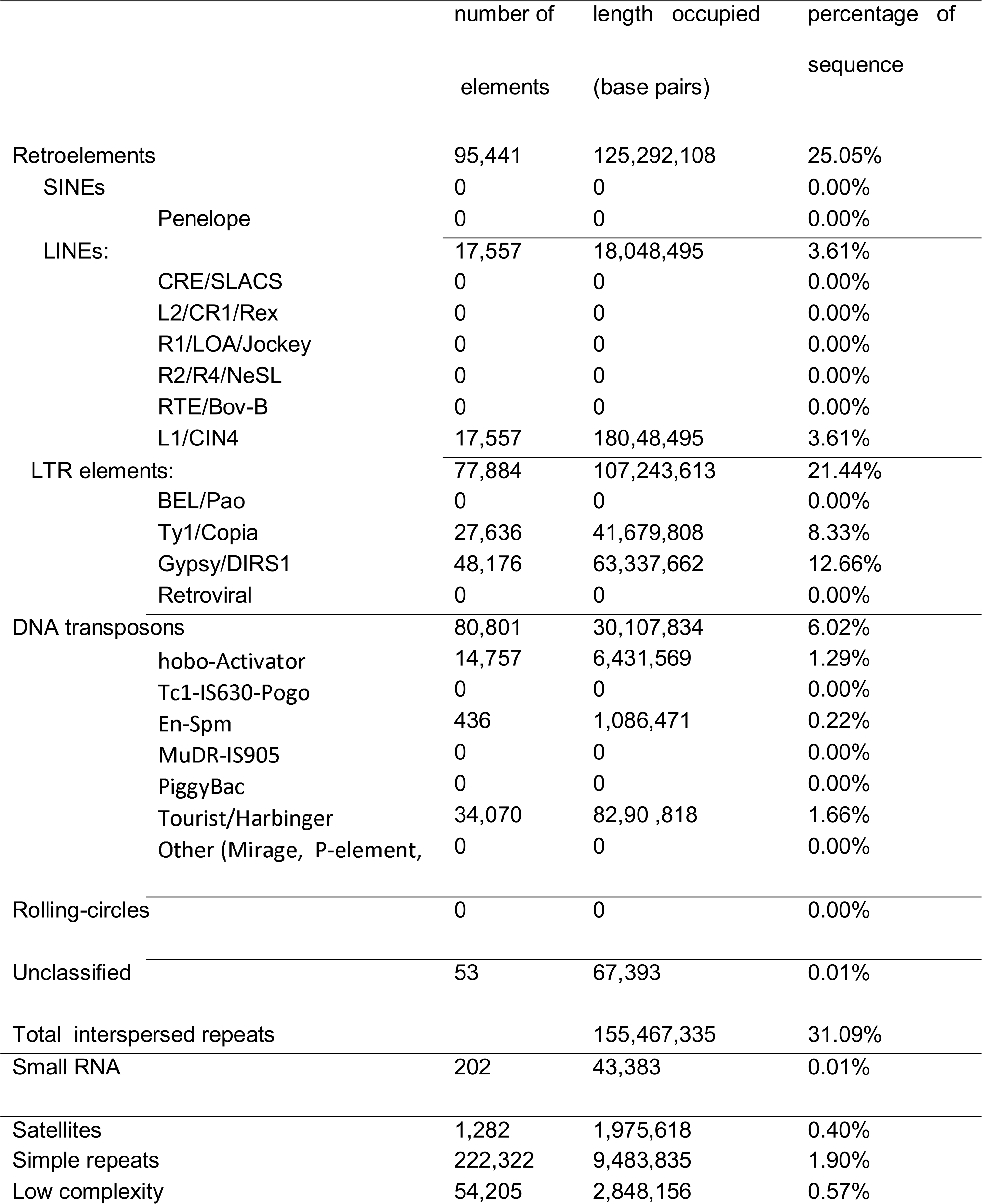
Repeated elements present in the *Vitis riparia* genome.

### Genotyping by Sequencing and genetic mapping

Two 96-plex GBS libraries (Keygene N.V. owns patents and patent applications protecting its Sequence Based Genotyping technologies) were constructed for the two parents (two replicates for each) and the 114 F1 plants of the cross CS ×□RGM. Raw reads were checked with FastQC^23^, demultiplexed with a custom script and cleaned with CutAdapt^24^. Cleaned reads were then mapped to the *V. riparia* RGM scaffolds previously obtained, the *V. vinifera* Cabernet Sauvignon contigs^12^ and *V. vinifera* PN40024 genome assemblies^4^ for SNP calling. Aligned on these genomes were performed using BWA^25^, SAMtools^26^ and Picard tools^27^ and SNP genotypes were detected with GATK^28^ using the *hardfilter* parameters^29^. In the variant call format (VCF) output file only sites with less than 20 % missing data and a minimum allele frequency (MAF)□≥□0.2 were retained. The SNP set was parsed into two data sets based on a pseudo-test cross mapping strategy^30^ using *major_minor* and *get_pseudo_test_cross* scripts from Hetmapps^31^. The segregation ratios of markers in the population were examined by Chi-square analysis. Markers with segregation ratios that differed from expected 1:1 at P<0.05 were classified as segregation distortion markers and discarded. The RGM and CS sets contain 1591 and 2359 SNPs respectively. Linkage groups (LGs) were determined using software JoinMap® 4.1^32 33^ and Rqtl^34^. LG were formed with a logarithm of odds (LOD) threshold of 6 and a maximum recombination frequency of 0.45. The 19 LGs that corresponded to the 19 chromosomes of grapevine were reconstructed and leaded to a total genetic map length of 2,268 cM and 2,514 cM for RGM and CS respectively.

### Pseudo-molecule construction

The PacBio / 10x Chromium Genomics hybrid scaffolding was organized into pseudo-molecules using GBS markers information from the CS x RGM genetic map. Scaffolds were anchored and oriented SNP using AllMaps^44^ with the *unequal weights2* parameters for a single run for the entire genome. Final pseudo-molecules were named according to *Vitis vinifera* PN40024 reference genome using SNP identification through SNP calling on this reference. Since PN40024 genome is the only one available who has been scaffolded into pseudo-molecules, collinearity with *V. riparia* was evaluated using D-GENIES^45^ and showed extremely high conservation along the 19 chromosomes of the species (Figure 1) even if the North American and Eurasian *Vitis* species diverged approximately 46.9 million years ago^46^.

**Fig. 1:**
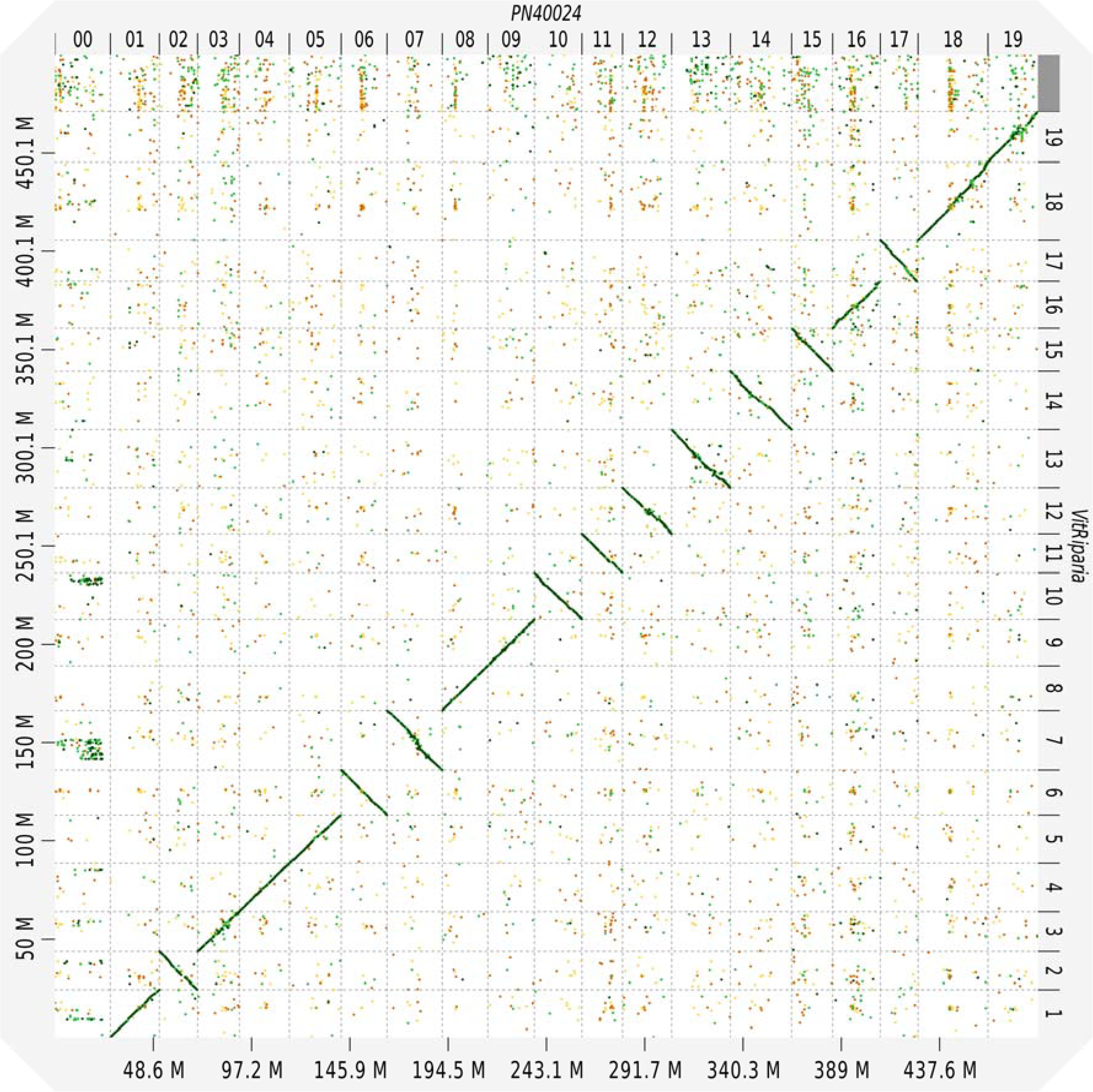
Comparison of *Vitis riparia* hybrid scaffolds with the reference PN40024 assembly. Hybrid scaffolds (Y-axis) were aligned to all 19 PN40024 chromosomes (X-axis) using D-GENIES and alignments were subsequently filtered for 1-on-1 alignments and rearrangements with a 20 Kbps length cutoff.

### Genome annotation and gene prediction

Consistent with observations that long reads sequencing technologies are a better solution for resolving repeat sequences, we found that known repetitive elements accounted for 170 Mb (33.94%) of the genome in *V. riparia*. This is a lower proportion among grape genomes when comparing published values to date. However, when comparisons are performed with the same analysis workflow and tools^47 48^, the percentages obtained between the two genotypes were in the same range (Online-only Table 1). Similar to other grape genomes, long terminal repeat (LTR) elements constituted the highest proportion of all repeated elements in *V. riparia*, (21.44%) with Copia and Gypsy families accounting for 8.33% and 12.66% respectively. The Long Interspersed Nuclear Elements (LINEs) and Miniature Inverted-repeat Transposable Elements (MITEs) represented 3.61% and 6.02% of the whole genome respectively.

After repeat masking, the genome was *ab initio* annotated using MAKER-P pipeline^49 50^, SNAP^51^ and Augustus^52^ gene finder with 3 rounds of Maker and an Augustus prediction. Structural annotation was then followed with an Interproscan functional annotation and putative gene function assignation using BLAST on UniProtKB. MAKER-P quality metrics with a threshold of AED<0.5 were chosen to retain the set of predicted genes. We finally generated a gene set of 37,207 protein-coding genes (11,434_AED<0.1; 8,638_0.1≤AED<0.2; 5,748_0.2≤AED<0.3; 5,418_0.3≤AED<0.4; 5,969_0.4≤AED<0.5) with 31,240 of them coupled with an evidence of protein function.

To facilitate genomic investigations for the community, a JBrowse Genome Browser^53^ was set up for *V. riparia* pseudo-molecules and is available from https://www6.bordeaux-aquitaine.inra.fr/egfv/.

## Code Availability

1. GBS demultiplexing

https://github.com/timflutre

2. Filters FASTQ files with CASAVA 2.20

fastq_illumina_filter --keep N -v -v -o good_reads.fq raw_reads.fastq

3. Cutadapt (regular 3’ adapter)

https://cutadapt.readthedocs.io/en/stable/guide.html

cutadapt -a AGATCGGAAGAGCGGTTCAGCAGGAATGCCGAG

-A AGATCGGAAGAGCGTCGTGTAGGGAAAGAGTGT

-G CTCGGCATTCCTGCTGAACCGCTCTTCCGATCT

-g ACACTCTTTCCCTACACGACGCTCTTCCGATCT -u7 -U7 -m10

4. Burrows-Wheeler Alignment – BWA-MEM

http://bio-bwa.sourceforge.net/bwa.shtml

bwa mem ref.fa read1.fastq.gz read2.fastq.gz > aligned.reads.sam with these options :

-M Mark shorter split hits as secondary (for Picard compatibility)

-R Complete read group header line with ‘\t’ used in STR to be converted to a TEB in the output SAM. An example is ‘@RG\tID:\tSM:\tPL:\tLB:’

5. Picard tools

https://broadinstitute.github.io/picard/

SortSam : java –jar picard.jar SortSam with these options: INPUT (BAM file), OUTPUT (BAM file), SORT_ORDER

MarkDuplicates : java –jar picard.jar MarkDuplicates with these options: INPUT (BAM file), OUTPUT (BAM file), METRIC_FILE (file)

BuildBamIndex : java –jar picard.jar BuildBamIndex with these options: INPUT (BAM file)

6. GATK tools

HaplotypeCaller : java –jar GenomeAnalysisTK.jar –T HaplotypeCaller –R ref.fasta –I file.bam –genotyping_mode DISCOVERY –drf DuplicateRead –emitRefConfidence GVCF –o file.g.vcf https://software.broadinstitute.org/gatk/documentation/tooldocs/3.8-0/org_broadinstitute_gatk_tools_walkers_haplotypecaller_HaplotypeCaller.php

CombineGVCFs : java –jar GenomeAnalysisTK.jar –T CombineGVCFs –R ref.fasta –drf DuplicateRead –G Standard –G AS_Standard --variant sample1 to sample’n’.g.vcf –o cohort_file.g.vcf https://software.broadinstitute.org/gatk/documentation/tooldocs/3.8-0/org_broadinstitute_gatk_tools_walkers_variantutils_CombineGVCFs.php

GenotypeGVCFs : java –jar GenomeAnalysisTK.jar –T GenotypeGVCFs –R ref.fasta –drf DuplicateRead –G Standard –G AS_Standard --variant cohort_file.g.vcf –o final_file.vcf https://software.broadinstitute.org/gatk/documentation/tooldocs/3.8-0/org_broadinstitute_gatk_tools_walkers_variantutils_GenotypeGVCFs.php

SelectVariants : java –jar GenomeAnalysisTK.jar –T SelectVariants –R ref.fasta –V final_file.vcf –selectType SNP –o file_snps.vcf https://software.broadinstitute.org/gatk/documentation/tooldocs/3.8-0/org_broadinstitute_gatk_tools_walkers_variantutils_SelectVariants.php

VariantFiltration : java –jar GenomeAnalysisTK –T VariantFiltration –R ref.fasta –V file_snps.vcf --filterExpression « QD < 2.0 || FS > 60.0 || MQ < 40.0 || MQRankSum < -12.5 || ReadPosRankSum < -8.0 » --filteredName «FILTER » -o filtered_snps.vcf https://software.broadinstitute.org/gatk/documentation/tooldocs/3.8-0/org_broadinstitute_gatk_tools_walkers_filters_VariantFiltration.php

7. VCF filtering

vcftools --vcf filtered_snps.vcf --remove-filtered-all --recode --out filteredFinal_snps.vcf

8. Falcon and Falcon_Unzip Assembly for SMRT sequencing https://github.com/PacificBiosciences/FALCON/wiki https://github.com/PacificBiosciences/FALCON_unzip/wiki

Main parameters: length_cutoff = 5000, length_cutoff_pr = 5000

pa_HPCdaligner_option = -v -dal128 -e0.70 -M40 -l2500 -k17 -h500 -w7 -s100

ovlp_HPCdaligner_option = -v -dal128 -M40 -k19 -h500 -e.96 -l1500 -s100

pa_DBsplit_option = -a -x500 -s200

ovlp_DBsplit_option = -s200

falcon_sense_option = --output_multi --output_dformat --min_idt 0.80 --min_cov 4 max_n_read 400 --n_core 16

falcon_sense_skip_contained = False

overlap_filtering_setting = --max_diff 120 --max_cov 120 --min_cov 4 --n_core 24

9. Purge Haplotigs

https://bitbucket.org/mroachawri/purge_haplotigs/src/master/ purge_haplotigs readhist -b aligned.bam -g genome.fasta

10. Supernova Assembly for 10x Chromium sequencing https://support.10xgenomics.com/de-novo-assembly/software/overview/latest/welcome Option *pseudohap2* style output

11. Scaffolding Falcon assembly with LINKS using Supernova outputs Assembly https://github.com/bcgsc/LINKS

LINKS -f.fa -s fileofname.fofn -b cns1-linked_draft -d 5000 -t 100 -k 19 -l 5 -a 0.3

LINKS -f.fa -s fileofname.fofn -b cns2-linked_draft -d 6000 -t 80 -k 19 -l 15 -a 0.3

LINKS -f.fa -s fileofname.fofn -b cns3-linked_draft -d 7000 -t 60 -k 19 -l 20 -a 0.3

LINKS -f.fa -s fileofname.fofn -b cns4-linked_draft -d 10000 -t 30 -k 19 -l 20 -a 0.3

LINKS -f.fa -s fileofname.fofn -b cns5-linked_draft -d 15000 -t 30 -k 19 -l 20 -a 0.3

LINKS -f.fa -s fileofname.fofn -b cns6-linked_draft -d 50000 -t 30 -k 19 -l 30 -a 0.3

LINKS -f.fa -s fileofname,fofn -b cns7-linked_draft -d 75000 -t 30 -k 19 -l 40 -a 0.3

12. Improving quality with PILON and Illumina sequencing https://github.com/broadinstitute/pilon/wiki/Requirements-&-Usage

13. Allmaps pseudomolecules scaffolding https://github.com/tanghaibao/jcvi/wiki/ALLMAPS

14. Assembly evaluation with BUSCO v3 https://busco.ezlab.org/

15. Vitis TE(s) Identification using RepeatMasker http://www.repeatmasker.org/

16. Annotation with MAKER_P pipeline, SNAP and Augustus gene finder http://www.yandell-lab.org/publications/pdf/maker_current_protocols.pdf https://bmcbioinformatics.biomedcentral.com/articles/10.1186/1471-2105-5-59 https://github.com/Gaius-Augustus/Augustus

DB (vitis) AND “Vitis”[porgn] from https://www.ncbi.nlm.nih.gov

EST DB (vitis) AND “Vitis”[porgn] from https://www.ncbi.nlm.nih.gov

- First run: rm_pass=0, est2genome=1 and protein2genome=1

gff3_merge -d master_datastore_index.log

maker2zff -c 0 -e 0 -o 0 -x 0.05 maker1.gff

fathom -categorize 1000 genome.ann genome.dna

fathom -export 1000 -plus uni.ann uni.dna

forge export.ann export.dna

hmm-assembler.pl RGM. > snap1.hmm

- Second run: rm_pass=1, est2genome=0, protein2genome=0, maker_gff=maker1.gff, snaphmm=snap1.hmm leading to a maker2.gff3 and a snap2.hmm files.

gff3_merge -d master_datastore_index.log

maker2zff -c 0 -e 0 -o 0 -x 0.05 maker2.gff

fathom -categorize 1000 genome.ann genome.dna

fathom -export 1000 -plus uni.ann uni.dna

forge export.ann export.dna

hmm-assembler.pl RGM. > snap2.hmm

Run Augustus:

zff2gff3.pl genome.ann | perl -plne ‘s/\t(\S+)$/\t\.\t$1/’ >genome.gff3

autoAug.pl --genome=../pilon2.fasta --species=RGM18 --cdna=sequence_est_ncbi.fasta – trainingset=genome.gff3 --singleCPU -v --useexisting

- Third run: rm_pass=1, est2genome=0, protein2genome=0, maker_gff=maker2.gff, snaphmm=snap2.hmm, augustus_species=RGM18 leading to a maker3.gff3, maker3.transcripts.fasta and maker3.proteins.fasta structural prediction.

gff3_merge -d master_datastore_index.log

fasta_merge -d master_datastore_index.log

17. Interproscan functional annotation and putative gene function assignation Download protein DB from http://www.uniprot.org

makeblastdb -in protein_db.fasta -input_type fasta -dbtype prot

blastp -db protein_db.fasta -query maker3.proteins.fasta -out maker3.proteins.blastp -evalue 0.000001 -outfmt 6 -max_hsps 1

maker_functional_gff protein_db.fasta maker3.proteins.blastp maker3.gff3 >> maker3.putative.gff3

maker_functional_fasta protein_db.fasta maker3.proteins.blastp maker3.proteins.fasta >> maker3.putative.proteins.fa

maker_functional_fasta protein_db.fasta maker3.proteins.blastp maker3.transcripts.fasta >> maker3.putative.transcripts.fa

Run Interproscan

interproscan.sh --iprlookup --goterms -f tsv -i maker3.putative.proteins.fa -pa -b RGM.annotated.proteins

18. Assembly validation using WBA mem

bwa mem -M -t 20 VitRiparia.fasta reads_pe.R1.fastq reads_pe.R2.fastq > aln_pe_reads.sam

samtools view -bS aln_pe_reads.sam -o aln_pe_reads.bam #| samtools sort - aln_reads.sorted.bam

samtools sort -o aln_pe_reads.sorted.bam aln_pe_reads.bam

bamtools stats -in aln_pe_reads.sorted.bam > bamstat_pe.reads

### Data records

The *V. riparia* genome project was deposited at NCBI under BioProject number PRJNA512170 and BioSample SAMN10662253. The DNA sequencing data from Illumina, PacBio and 10x Genomics have been deposited in the Sequence Read Archive (SRA) database under accession numbers SRX5189632 to SRX5189680. This Whole Genome Shotgun project has been deposited at DDBJ/ENA/GenBank under the accession SJAQ00000000. The version described in this paper is version SJAQ01000000. Genetic mapping data and structural and functional annotation file of the *Vitis riparia* assembly are available on figshare (https://figshare.com/s/0a52d4408214e9f1e280).

### Technical validation

To evaluate the accuracy and completeness of the *V. riparia* assembly, genome features were compared to those of *V. vinifera* (Table 2). We found that both contig and scaffold N50 lengths of *Vitis riparia* reached considerable continuity. The Guanine-Cytosine content (GC = 34.32 %) was similar to those of *V. vinifera* Chardonnay (34.43%).

To further assess the accuracy of the *V. riparia* genome assembly, the NGS-based short reads from whole-genome sequencing data were also aligned against the genome assembly using BWA mem^54^. We found that 98.4% of the reads were reliably aligned to the genome assembly, and 95.8% of the reads were properly aligned to the genome with their mates. Paired-end reads data were not used during the contig assembly, thus the high alignment ratio demonstrated the high quality of contig assembly.

The assembled genome was also subjected to Benchmarking Universal Single-Copy Orthologs^55^, which quantitatively assesses genome completeness using evolutionarily informed expectations of gene content from near-universal single-copy orthologs, using the genes in the embryophyta release 9 dataset (embryophyta.odb9). The BUSCO results showed that 96.5% of conserved BUSCO proteins were detected in the *V. riparia* assembly, including 1.1% of fragment BUSCO proteins (Table 2). Overall, these metrics compare well with other recently published grape genomes, providing a high quality genome sequences for the following functional investigations.

## Conflicts of interests

The authors declare that they have no competing interests.

### Acknowledgements

We thank Dr. Nathalie Ollat and Dr. Gregory Gambetta for critical review of the manuscript. We thank the INRA GeT-PlaGE platform (http://get.genotoul.fr) for sequencing the genome, the INRA Genotoul bioinformatics platform (http://genotoul.toulouse.inra.fr) for providing computational resources and support.

We thank Dr. Timothée Flutre and Amandine Launay for their help in coordinating the FruitSelGen project and in acquiring GBS data.

This work was supported by a grant overseen by the French National Research Agency (ANR) as part of the ANR-09-GENM-024 “Vitsec” program and by the University of Bordeaux.

## Author contributions

Author P.-F. B. conceived the project. N. G. and P.-F. B. assembled the genomes, performed the genome annotation and downstream analyses. B. R. performed GBS analysis and genetic mapping. P.-F. B. wrote the paper. All authors read, edited and approved the final manuscript.

## Notes

https://figshare.com/s/0a52d4408214e9f1e280

https://www.ncbi.nlm.nih.gov/assembly/GCA_004353265.1

https://www.ncbi.nlm.nih.gov/bioproject/PRJNA512170

## References

1. Peterson, D.G. & Arick, M. Sequencing Plant Genomes. (Progress in Botany. Springer, Berlin, Heidelberg 2018).

2. Nguyen, K.L., Grondin, A., Courtois, B., & Gantet, P. Next-generation sequencing accelerates crop gene discovery. Trends Plant Sci 24, 263–274 (2018).

3. Scheben, A., Yuan, Y. & Edwards, D. Advances in genomics for adapting crops to climate change. Curr. Plant Biol. 6, 2–10 (2016).

4. Jaillon, O. et al. The grapevine genome sequence suggests ancestral hexaploidization in major angiosperm phyla. Nature 449, 463–467 (2007).

5. Velasco, R. et al. A High Quality Draft Consensus Sequence of the Genome of a Heterozygous Grapevine Variety. PLoS ONE 2:e1326 (2007).

6. Alkan, C., Sajjadian, S. & Eichler, E.E. Limitations of next-generation genome sequence assembly. Nat. Methods 8,61–65 (2010).

7. Da Silva, C. et al. The high polyphenol content of grapevine cultivar Tannat berries is conferred primarily by genes that are not shared with the reference genome. Plant Cell 25, 4777–4788 (2013).

8. Di Genova, A. et al. Whole genome comparison between table and wine grapes reveals a comprehensive catalog of structural variants. BMC Plant Biol 14, 7 (2014).

9. Jiao, W.B. & Schneeberger, K. The impact of third generation genomic technologies on plant genome assembly. Curr. Opin. Plant Biol. 36, 64–70 (2017).

10. Li, C., Lin, F., An, D., Wang, W. & Huang, R. Genome Sequencing and Assembly by Long Reads in Plants. Genes 9, 6 (2018).

11. Treangen, T.J. & Salzberg, S.L. Repetitive DNA and next-generation sequencing: computational challenges and solutions. Nat. Rev. Genet. 13, 36–46 (2012).

12. Chin, C.S. et al. Phased diploid genome assembly with single-molecule real-time sequencing. Nat. Methods 13, 1050–1054 (2016).

13. Roach, M. J., Schmidt, S. A., & Borneman, A. R. Purge Haplotigs: allelic contig reassignment for third-gen diploid genome assemblies. BMC bioinformatics 19, 460 (2018).

14. Vinson, J.P. et al. Assembly of polymorphic genomes: algorithms and application to *Ciona savignyi*. Genome Research 15, 1127–35 (2005).

15. Roach, M.J. et al. Population sequencing reveals clonal diversity and ancestral inbreeding in the grapevine cultivar Chardonnay. PLoS Genet 14:e1007807.(2018)

16. Zhou, Y.S. et al. Structural variants, clonal propagation, and genome evolution in grapevine (*Vitis vinifera*) bioRxiv 508119; doi: https://doi.org/10.1101/508119 (2018).

17. Laucou V. et al. Extended diversity analysis of cultivated grapevine Vitis vinifera with 10K genome-wide SNPs. PLoS One 13:e0192540 (2018).

18. Myles, S. et al. Genetic structure and domestication history of the grape. Proc Natl Acad Sci U S A 108, 3530–3535 (2011).

19. Migicovsky, Z. et al. Genomic ancestry estimation quantifies use of wild species in grape breeding. BMC Genomics 17, 478 (2016).

20. FAO Commission on genetic resources for food and agriculture assessment. The state of the world’s biodiversity for food and agriculture (2019).

21. Marguerit, E. et al. Genetic dissection of sex determinism, inflorescence morphology and downy mildew resistance in grapevine. Theor. Appl. Genet. 118, 1261–1278 (2009).

22. Elshire, R. J. et al. A robust, simple genotyping-by-sequencing (GBS) approach for high diversity species. PLoS ONE 6:e19379 (2011).

23. Andrews, S. Fastqc: a quality control tool for high throughput sequence data, http://www.bioinformatics.babraham.ac.uk/projects/fastqc (2010).

24. Martin, M.. Cutadapt removes adapter sequences from high-throughput sequencing reads. EMBnet.journal 17, 10–12 (2011).

25. Li, H. et al. The Sequence Alignment/Map format and SAMtools. Bioinformatics 25, 2078–2079 (2009).

26. Li, H. & Durbin, R. Fast and accurate long-read alignment with Burrows–Wheeler transform. Bioinformatics 26, 589–595 (2010).

27. Picard Tools - By Broad Institute. Available from: http://broadinstitute.github.io/picard.

28. DePristo M.A. et al. A framework for variation discovery and genotyping using next-generation DNA sequencing data. Nat Genet 43, 491–498 (2011).

29. Van der Auwera, G.A., et al. From FastQ Data to High-Confidence Variant Calls: The Genome Analysis Toolkit Best Practices Pipeline. Current Protocols in Bioinformatics 43, 1–11 (2013).

30. Grattapaglia, D., Bertolucci, F.L.G & Sederoff, R. Genetic mapping of QTLs controlling vegetative propagation in *Eucalyptus grandis* and *E. urophylla* using a pseudo-testcross mapping strategy and RAPD markers. Theor. Appl. Genet. 90, 933–947 (1995).

31. Hyma, K.E. et al. Heterozygous mapping strategy (HetMappS) for high resolution Genotyping-By-Sequencing markers: a case study in grapevine. PLoS One 10: e0134880 (2015).

32. Stam, P. & Van Ooijen J.W. JOINMAP version 2.0: software for the calculation of genetic linkage maps (1995).

33. Van Ooijen, J.W. JoinMap® 4.0, Software for the calculation of genetic linkage maps in experimental populations. Kyazma B.V. Wageningen, Netherlands (2006)

34. Broman, K.W., Wu, H., Sen, S., & Churchill G.A. R/qtl: QTL mapping in experimental crosses. Bioinformatics 19, 889–890 (2003).

35. Lodhi, M.A. & Reisch, B.I. Nuclear DNA content of *Vitis* species, cultivars, and other genera of the Vitaceae. Theor. Appl. Genet. 90, 11–16 (1995).

36. Liu, B., Shi, Y., Yuan, Y., Hu, X., Zhang, H., Li, N., Li, Z., Chen, Y., Mu, D., and Fan, W.. Estimation of genomic characteristics by analyzing k-mer frequency in de novo genome projects. arXiv preprint arXiv:1308 (2012).

37. Marcais, G. & Kingsford, K. A fast, lock-free approach for efficient parallel counting of occurrences of k-mers. Bioinformatics 27, 764–770 (2011).

38. PacBio FALCON GitHub page https://github.com/PacificBiosciences/FALCON, Accessed 18 Mar 2018.

39. Pacific Biosciences, SMRT tools. https://www.pacb.com/wp-content/uploads/SMRT-Tools-Reference-Guide-v4.0.0.pdf.

40. Chaisson, M.J. & Tesler, G. Mapping single molecule sequencing reads using basic local alignment with successive refinement (BLASR): application and theory. BMC Bioinformatics 13, 238 (2012).

41. Walker, B.J. et al. Pilon: an integrated tool for comprehensive microbial variant detection and genome assembly improvement. PLoS One 9:e112963 (2014).

42. Weisenfeld, N.I., Kumar, V., Shah, P., Church, D.M. & Jaffe D.B. Direct determination of diploid genome sequences. Genome research 27, 757–767 (2017).

43. Warren, R.L. et al. LINKS: Scalable, alignment-free scaffolding of draft genomes with long reads. Gigascience. 4, 35 (2015).

44. Tang, H. et al. ALLMAPS: robust scaffold ordering based on multiple maps. Genome Biol. 16. 3–10 (2015).

45. Cabanettes, F. & Klopp, C. D-GENIES: dot plot large genomes in an interactive, efficient and simple way. PeerJ 6:e4958 (2018).

46. Ma, Z.Y. et al. Phylogenomics, biogeography, and adaptive radiation of grapes. Molecular phylogenetics and evolution 129, 258–267 (2018).

47. Tarailo-Graovac, M. & Chen, N. Using RepeatMasker to identify repetitive elements in genomic sequences. Curr Protoc Bioinformatics 25, 4–10 (2009).

48. Bao, W., Kojima, K.K. & Kohany, O. Repbase Update, a database of repetitive elements in eukaryotic genomes. Mobile DNA 6:11 (2015).

49. Campbell, M.S., Holt, C., Moore, B. & Yandell, M. Genome Annotation and Curation Using MAKER and MAKER-P. Curr Protoc Bioinformatics 48, 1–39 (2014).

50. Campbell, M.S. et al. MAKER-P: a tool kit for the rapid creation, management, and quality control of plant genome annotations. Plant Physiol 164, 513–24. (2014).

51. Korf, I. Gene finding in novel genomes. BMC Bioinformatics 5:59. (2004).

52. Stanke, M. et al. AUGUSTUS: a web server for gene finding in eukaryotes. Nucleic Acids Research 32, 309–312 (2004).

53. Buels, R. et al. JBrowse: a dynamic web platform for genome visualization and analysis. Genome Biology 12, 17–66 (2016).

54. Li H. Aligning sequence reads, clone sequences and assembly contigs with BWA-MEM. arXiv:1303.3997 1. (2013)

55. Simão, F.A., Waterhouse, R.M., Ioannidis, P., Kriventseva, E.V. & Zdobnov, E.M. BUSCO: assessing genome assembly and annotation completeness with single-copy orthologs. Bioinformatics 31, 3210–3212 (2015).

## Data Citations

1. NCBI Sequence Read Archive https://www.ncbi.nlm.nih.gov/bioproject/PRJNA512170 (2018)

2. NCBI Assembly https://www.ncbi.nlm.nih.gov/assembly/GCA_004353265.1 (2019)

3. Girollet, N., Rubio, B. and Bert P.-F. De novo phased assembly of the Vitis riparia grape genome. figshare https://figshare.com/s/0a52d4408214e9f1e280 (2019)

